# Silencing neuroinflammatory reactive astrocyte activating factors ameliorates disease outcomes in perinatal white matter injury

**DOI:** 10.1101/2022.12.19.521083

**Authors:** Patricia Renz, Daniel Surbek, Valérie Haesler, Vera Tscherrig, Eric J Huang, Manideep Chavali, Shane Liddelow, David Rowitch, Andreina Schoeberlein, Amanda Brosius Lutz

## Abstract

The role of reactive astrocytes in perinatal white matter injury (WMI) is unclear. In a mouse model of WMI, we provide evidence that impairing the formation of a *C3*-expressing neuroinflammatory reactive astrocyte sub-state rescues myelination and behavioral deficits. We further demonstrate the presence of *C3*-expressing reactive astrocytes in human WMI. Our data point to these cells as putative drivers of myelination failure in WMI and a potentially promising therapeutic target.

## Main Text

Perinatal white matter injury is the most common cause of long-term neurological morbidity in infants born preterm [1]. Resulting from a series of hypoxic-ischemic insults to the developing brain during a key window of oligodendrocyte cell lineage vulnerability between the 23^rd^ and 32^nd^ week of human gestation, WMI is characterized histologically by morphological ‘reactivity’ changes in astrocytes and microglia and by myelination defects in periventricular white matter [1–3]. Clinically, patients present with a broad spectrum of motor and cognitive deficits [1]. Treatment options are currently extremely limited, in large part hampered by a limited understanding of underlying disease pathophysiology [1, 2, 4–6].

Silencing the expression of activated microglia-derived cytokines *Tnf, Il1a*, and *C1q* improves disease outcomes in multiple neurodegenerative disease models in adult mice [7–11]. Similarly, in large and small animal models of WMI, in vivo TNFα blockade and attenuation of microglia-mediated neuroinflammation, respectively, have been shown to improve myelination defects [12–14]. Based on these findings, we aimed to examine the effect of silencing *Tnf, Il1a*, and *C1q* on WMI disease outcomes during early development. Using an established rodent model of perinatal WMI recently published by our group, we quantified myelination in the corpus callosum at postnatal day 11 (9 days post injury, dpi) in *Tnf, Il1a, C1q* triple knockout (TKO) mice in comparison with wildtype (WT) C57Bl/6 mice (Figure 1A) [15]. We observed the expected myelination defects in injured wildtype mice. In contrast, myelination was not significantly impaired in injured TKO mice (Figure 1B). Myelination in the corpus callosum was not significantly different between uninjured WT and uninjured TKO mice (Supplementary Figure 1).

**Figure 1.**
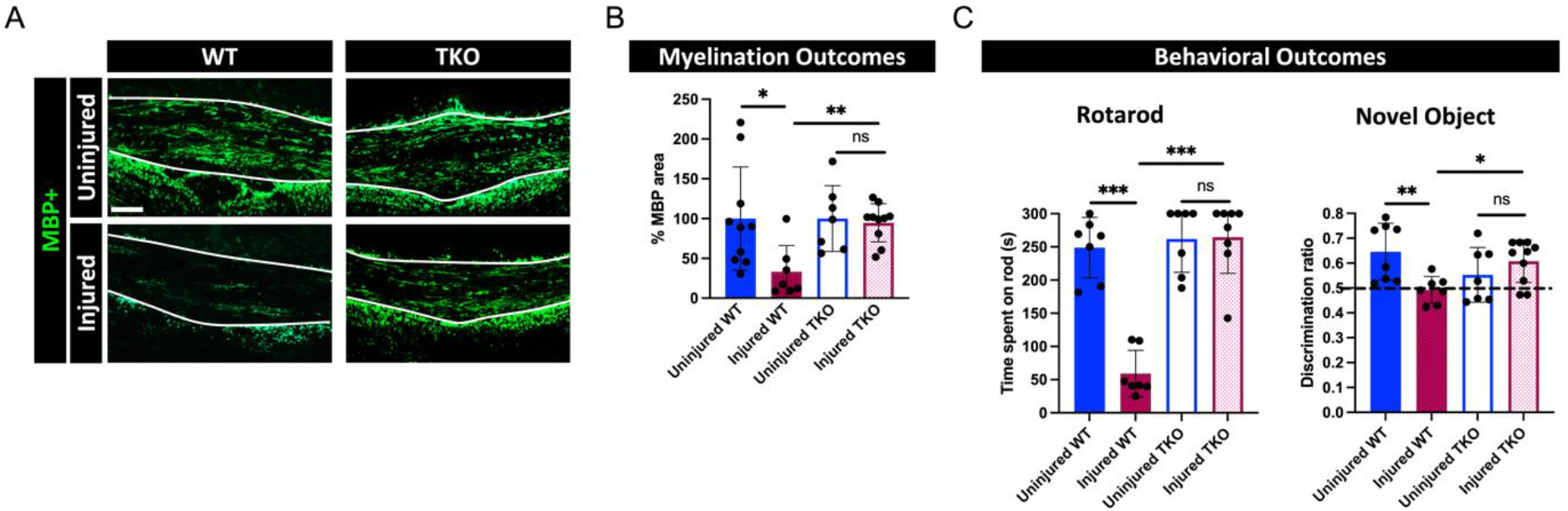
Quantification of myelination defects and behavioral outcomes after perinatal white matter injury in *Tnf, Il1a, C1q* triple knockout (TKO) and wildtype (WT) C57Bl/6mice. (A) Representative images of myelin basic protein immunofluorescence (IF) (green) in the medial corpus callosum of uninjured and injured WT and TKO mice at 9 days post injury. Scale bar: 100 μm. (B) Quantification of MBP immunofluorescence in the medial corpus callosum of uninjured and injured WT and TKO mice. Percentage areas were normalized to the mean % area of uninjured mice. Uninjured WT (n = 10), injured WT (n = 7), uninjured TKO (n = 7), injured TKO (n = 10). (C) Quantification of rotarod (left) and novel object recognition (right) test performance. Uninjured WT (n = 7), injured WT (n = 7), uninjured TKO (n = 7), injured TKO (n = 8). Data are presented as mean ± SD. ns not significant, \**p* <0.05, \*\**p* < 0.01, \*\*\**p* < 0.001.

Our rodent model of perinatal white matter injury results in functional deficits that mirror the sensorimotor and cognitive disabilities observed in humans affected by WMI [15]. Given reduced myelination defects in TKO mice, we also aimed to investigate whether the absence of TNFα, IL1α, and C1q could rescue behavioral outcomes in our WMI model. Using rotarod and novel object recognition testing at 28 dpi, we observed the expected deficits in motor performance and recognition memory in injured compared with uninjured WT mice. In contrast to injured WT mice, injured TKO mice were able to perform both behavioral tasks as well as uninjured mice (Figure 1C-D).

Evidence from multiple groups indicates that microglial-derived cytokines TNFα, IL1α, and C1q induce the formation of a complement component 3 (*C3*)-expressing neuroinflammatory reactive astrocyte sub-state reported to be toxic to neurons and mature oligodendrocytes, and to block OPC-to-oligodendrocyte differentiation in culture [10, 16–19]. The role of reactive astrocytes in WMI disease pathogenesis is currently unclear [20–22]. Our results in TKO mice in our mouse model of WMI suggest that microglia-induced changes in astrocyte reactivity may play an important role in production of the myelination defects and functional deficits that characterize WMI. To test this hypothesis, we next examined whether *C3*-expressing astrocytes form in the corpus callosum in our rodent model of acute perinatal WMI. In situ hybridization for *C3* in WT mice at 1 dpi revealed that the percentage of astrocytes expressing *C3* is significantly higher in the injured corpus callosum compared with the uninjured corpus callosum (Figure 2A, B, Supplementary Figure 2A). Approximately one-third of astrocytes in the injured WT corpus callosum express *C3* at 1 dpi compared with approximately 3% of astrocytes in the uninjured WT corpus callosum (Figure 2B). The formation of these *C3*- expressing astrocytes was abrogated in TKO mice (Figure 2B). When we examined later timepoints after injury, we found that the proportion of astrocytes expressing *C3* remained significantly elevated in the injured relative to the uninjured corpus callosum at both 2 dpi and 9 dpi in WT mice (Figure 2C).

**Figure 2.**
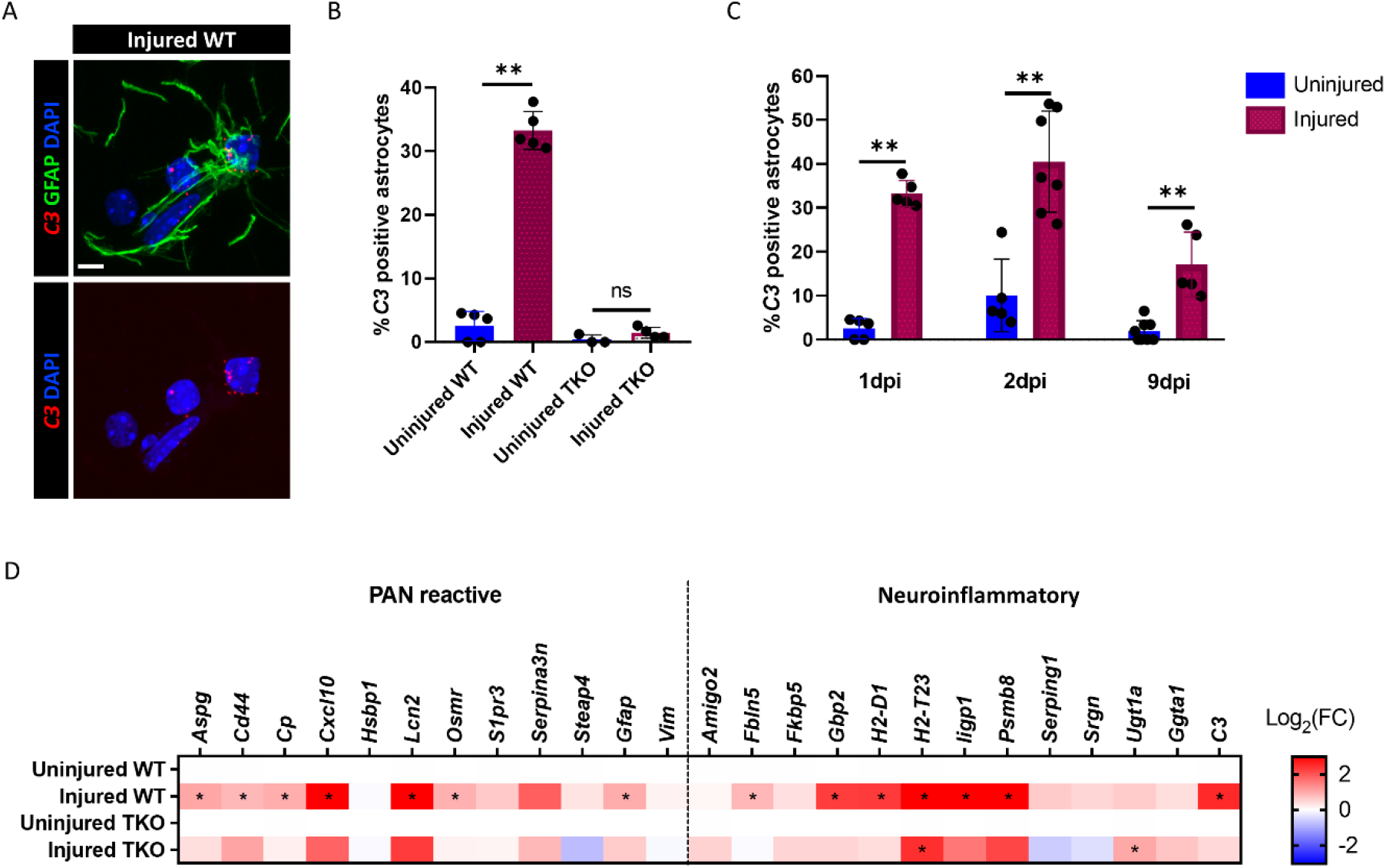
Formation of neuroinflammatory (C3+) astrocytes in rodent model of perinatal white matter injury in *Tnf, Il1a, C1q* triple knockout (TKO) and wildtype (WT) C57Bl/6 mice. (A) In situ hybridization for neuroinflammatory reactive astrocyte marker *C3* (red) and immunohistochemistry for GFAP (green) in the mouse corpus callosum of an injured WT brain. Scale bar: 10 μm. (B) Quantification of in situ hybridization for astrocyte markers *Aldh1l1* and *Gfap* (probe mix) and neuroinflammatory astrocyte marker *C3*. The graph depicts the percentage of astrocytes expressing *C3* in the corpus callosum at 1 dpi in WT (n = 5) and TKO (n = 4) mice. (C) Time course of *C3*-expressing reactive astrocyte formation in injured and uninjured WT mice. Cells were quantified using in situ hybridization as in (B). The graph depicts the percentage of astrocytes expressing *C3* in the corpus callosum in uninjured (1 dpi: n = 5; 2 dpi: n = 5; 9 dpi: n = 10) and injured (1 dpi: n = 5; 2 dpi: n = 7; 9 dpi: n = 7) WT mice. (D) Microfluidic qRT-PCR for a panel of genes broadly upregulated in many reactive astrocyte sub-states (“PAN reactive”), and genes characteristic of the neurotoxic astrocyte response to neuroinflammation (“neuroinflammatory”). Heatmap compares the mean fold change (FC) of “PAN reactive” and neurotoxic neuroinflammatory gene transcripts from astrocytes isolated from the cortex and white matter of uninjured and injured WT (n = 6) or TKO (n = 4) mice. Exact *p*-values for the heatmap are listed in Supplementary Table 2. Data are presented as mean ± SD. ns not significant, \**p* <0.05, \*\**p* < 0.01.

To more deeply understand changes in astrocyte reactivity induced by our WMI model, we examined the expression of genes broadly associated with astrocyte reactivity [16, 23]. Using microfluidics qRT-PCR of FACS-purified astrocytes isolated from the cortex and underlying white matter of mice at 1 dpi, we demonstrated significant upregulation of numerous reactivity genes that are representative of many reactive astrocyte sub-states in astrocytes of injured WT mice [17, 23, 24]. We refer these transcripts as ‘PAN reactive’ genes and they include *Aspg, Cd44, Cp, Cxcl10, Lcn2, Serpina3n*, and *Gfap* (Figure 2D, Supplementary Figure 2B).

In addition to the expression of *C3*, exposure to microglial-derived cytokines TNFα, IL1α, and C1q in response to systemic inflammation induces the upregulation of additional astrocyte gene transcripts at the bulk RNA sequencing level characteristic of the neurotoxic astrocyte response to neuroinflammation [16, 23, 24]. We investigated whether these additional transcriptional changes were occurring in astrocytes in our WMI model. Using qRT-PCR and FACS purification of astrocytes from injured WT C57Bl/6 mice, we demonstrated a significant upregulation of multiple ‘neuroinflammatory’ astrocyte gene transcripts (Figure 2D).

When we performed the same qRT-PCR analysis on astrocytes isolated from TKO mice, nearly all (13/14) of the significant changes in ‘PAN reactive’ and ‘neuroinflammatory’ gene expression were no longer observed. This finding demonstrates that silencing the microglia-derived expression of cytokines *Tnf, Il1a*, and *C1q* blocks astrocyte conversion towards the previously described neurotoxic, neuroinflammatory state (Figure 2D).

While our ongoing experiments aim to further define the molecular identity of neuroinflammatory reactive astrocytes in WMI and to solidify their role as drivers of disease pathology, the ability to rescue myelination and behavioral outcomes in TKO mice lacking *C3*-expressing astrocytes suggests that these cells may be promising therapeutic targets in this disease.

Although our rodent WMI model involves a hypoxic insult in addition to an inflammatory insult, at the cell population level, we did not observe significant upregulation of the gene expression pattern formerly attributed to “A2” reactivity and initially described in association with ischemic injury (Supplementary Figure 2C) [16, 23]. Without examining reactive astrocyte identity at the single cell level, however, the concomitant existence of astrocytes polarized in terms of gene expression in this direction cannot be excluded, and it is a response we would expect given recent investigations into reactivity heterogeneity by our groups and others [24–26].

To determine whether the formation of *C3*-expressing neuroinflammatory reactive astrocytes is also a feature of human perinatal WMI, we obtained human post-mortem brain tissue from infants affected by perinatal WMI and control cases matched as closely as possible based on gestational age at birth, age at time of death, and brain region available. The diagnosis of diffuse WMI was determined post-mortem by a neuropathologist. Both of the WMI cases that we examined involved preterm birth during the window of heightened pre-oligodendrocyte vulnerability between approximately 23 and 32 weeks gestation (Figure 3A). The infant we refer to as Case 1 survived for only 12 days after birth, while the other infant (Case 2) survived until 8 months of age, allowing us to assess the presence of *C3*-expressing astrocytes at both a short and a long time interval after injury (Figure 3A). The tissue obtained was from the globus pallidus and internal capsule region (Case 1) and from the cingulate cortex and corpus callosum region (Case 2). Using *C3* (ISH) and GFAP (IHC), we examined the proportion of GFAP positive astrocytes that also express *C3* in the white matter regions of this human tissue as well as in the same anatomical regions of the matched control cases. Over two-thirds (67%) of astrocytes in the internal capsule region of Case 1 expressed *C3* in comparison with 5% in the control brain (Figure 3B-C, D). In Case 2, one-third (33%) of astrocytes expressed *C3* in comparison with 5% in the control brain (Figure 3E). These results confirm the relevance of the formation of *C3*-expressing reactive astrocytes to human WMI and suggest that at least a subpopulation of these cells persists up to several months after injury.

**Figure 3.**
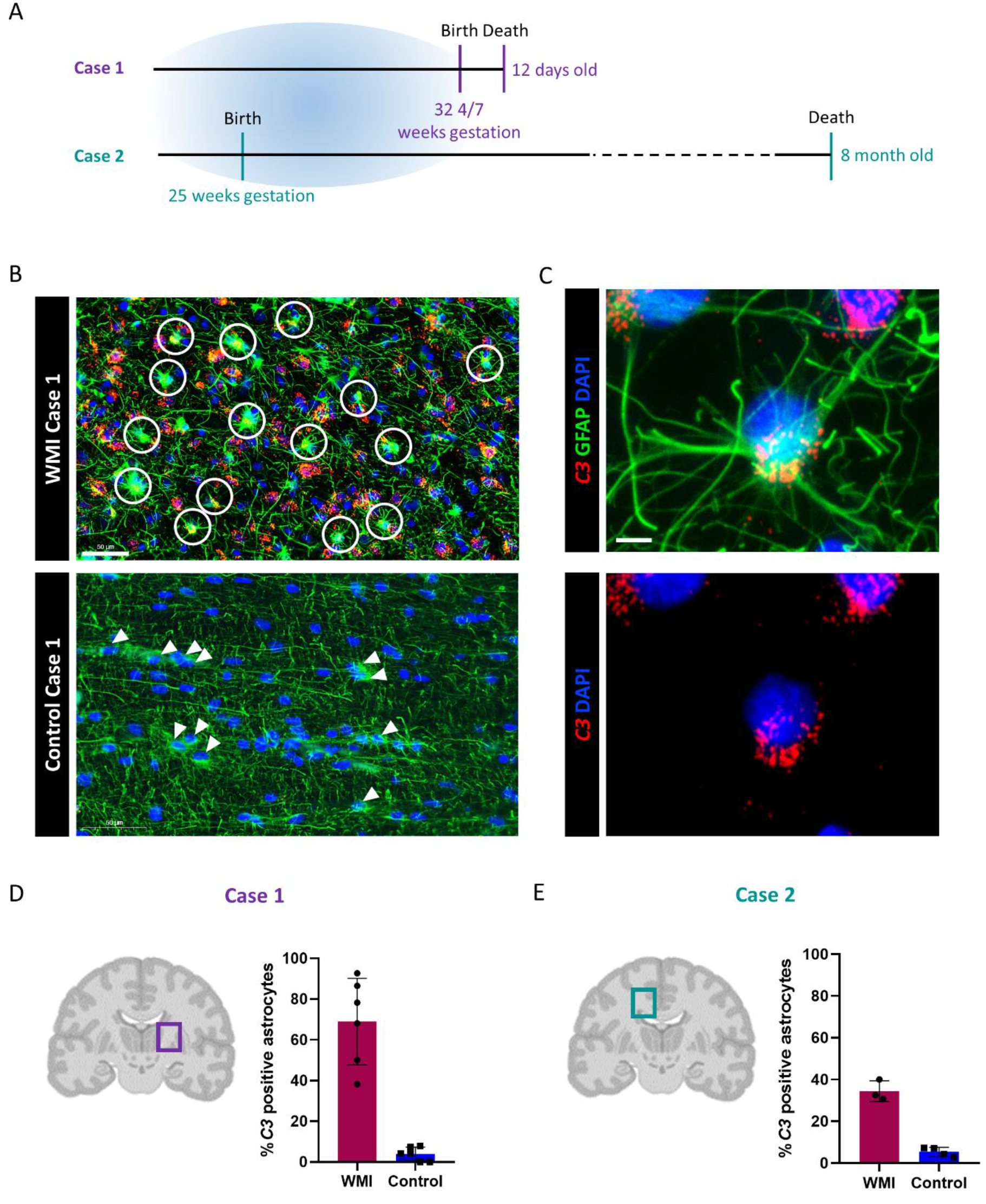
Formation of *C3*-expressing astrocytes in the human brain of perinatal white matter injury. (A) Timeline of gestational age at birth and postnatal age at time of death for white matter injury case 1 (purple) and case 2 (green). The blue area indicates the time window of pre-oligodendrocyte vulnerability. (B) Representative images of the human post-mortem tissue of the WMI Case 1 and Control Case 1. White circles indicate astrocytes co-expressing *C3* (red) and GFAP (green) in WMI Case 1. Arrowheads indicate astrocytes (green) in Control Case 1. Scale bar: 50 μm. (C) Confocal image of in situ hybridization for neurotoxic neuroinflammatory reactive astrocyte marker probe *C3* (red) and immunohistochemistry for GFAP (green) in the human WMI Case 1. Scale bar: 10 μm. (D-E) Coronal brain sections indicating the brain region used for quantifying the percentage of astrocytes expressing marker *C3* in Case 1 (D) and Case 2 (E) in WMI and matched control cases. Data points represent the percent of astrocytes expressing C3 in single 0.5 mm^2^ circles spaced regularly throughout the anatomical region of interest.

Consistent with the notion of reactive astrocyte heterogeneity, only a subpopulation of astrocytes in the white matter regions we examined (approximately 30% of astrocytes in the mouse corpus callosum, approximately 60% of astrocytes in the human internal capsule) expressed *C3*. This proportion is comparable to the fraction of astrocytes expressing *C3* in other disease models in which the abrogation of neuroinflammatory astrocytes has been shown to improve disease outcomes; in these studies C3 positive astrocytes accounted for approximately 30-60% of astrocytes in the region of pathology, but nearly none of the astrocytes distant from pathology [7–9].

Elaborate work in recent months has underlined the heterogeneity of astrocytes in health and disease [24, 25, 27, 28]. In light of these studies, the neuroinflammatory astrocyte conversion referred to in this manuscript is almost certainly a composite of multiple reactive astrocyte sub-states with inter- and intraregional diversity. Fully understanding the landscape of astrocyte reactivity and its functional implications in any disease state ultimately requires gene expression analysis at single cell resolution, data on the spatial coordinates of gene expression, and extensive experimental studies using in vitro modeling and manipulation of gene expression to uncover the functional consequences of these transcriptomically-defined reactive sub-states. Using these tools, it will be fascinating to uncover unique properties of astrocyte reactivity in response to the combined inflammatory/hypoxic injury of WMI and to elucidate differences in astrocyte responses to injury in the perinatal versus adult brain.

In summary, our findings show that impairing the formation of a *C3*-expressing neuroinflammatory reactive astrocyte sub-state by blocking expression of the microglial-derived cytokines TNFα, IL-1α, and C1q rescues WMI-associated defects in myelination as well as behavioral deficits. These results provide a basis for novel therapeutic approaches to preventing long-term neurodevelopmental deficits in infants born preterm.

## Methods

### Animals

All animal procedures were approved by the Veterinary Department of the Canton of Bern, Switzerland (reference number: BE19/85) and kept under standard housing conditions. We used C57BL/6 (WT; B6.Tg(Aldh1l1–eGFP)<OFC789Gsat>/Mmucd) and triple knockout (TKO; B6.Cg-Il1a<tm1Yiw>Tnf<tm1Gkl>C1qa<tm1Barr>Tg(Aldh1l1–eGFP)<OFC789Gsat>/Barr) mice. TKO mice were a gift from SAL.

### Mouse model of perinatal WMI

The two-hit acute diffuse perinatal white matter injury was induced, according to Renz et al. [15]. Briefly, following random assignment to an injured or an uninjured group. In the injury group, mouse pups were injected subcutaneously between the skapulae with 2 mg/kg body weight (BW) lipopolysaccharide (LPS; Escherichia coli strain O55:B5, Sigma Aldrich) or the equivalent volume of saline on postnatal day (P)2, placed on an isolette and submitted to a hypoxic insult (8% O_2_/92% N_2_, 3 l/min) for 25 min 6 hours after LPS injection. Uninjured mice received an equivalent volume of saline and were placed for 25 min 6 hours after saline injection in a temperature-controlled, normoxic environment.

### Immunohistochemistry on mouse tissue

At 9 days after injury (9 dpi), injured and uninjured WT and TKO mice were terminally anesthetized and perfused transcardially with phosphate-buffered saline (PBS) followed by 4% paraformaldehyde (PFA). Brains were removed and fixed in 4% PFA for 22 hours at 4°C. After fixation, all brains were embedded in paraffin and 6 μm sections in the coronal plane at the level of the hippocampus were cut using a microtome. Prior to staining, sections were deparaffinized and rehydrated. After antigen retrieval in 0.1 M sodium citrate buffer in a pressure cooker for 12 min, sections were washed in 1x Tris-buffered saline (TBS) with 0.1% Tween 20 (Sigma Aldrich) and blocked for 1 h at room temperature (RT) with 10% goat serum, 1% bovine serum albumin (BSA, Sigma Aldrich) in TBS. Sections were then incubated overnight at 4°C with the polyclonal rabbit anti- myelin basic protein (MBP) antibody (1:200, ab40390, Abcam) diluted in blocking buffer. Sections were next washed three times in 1x TBS and incubated with the secondary antibody (Alexa fluor^®^ 488-conjugated goat anti-rabbit IgG, Thermo Fisher Scientific) for 1 hour at RT in the dark. Sections were counterstained with DAPI (Sigma Aldrich).

### RNAscope^®^ in situ hybridization

RNAscope^®^ fluorescent in situ hybridization was performed on paraffin embedded brain sections. Mice pups 1 day post injury (1 dpi) and 2 days post injury (2dpi) were euthanized using rapid decapitation. Brains were removed and directly fixed in 4% paraformaldehyde (PFA) for 20 h at 4°C and embedded in paraffin. Nine dpi tissue was obtained as described in the immunohistochemistry methods. RNAscope^®^ Multiplex Fluorescent Reagent Kit v2 (Cat. Nr. 323110, Advanced Cell Diagnostics (ACDbio)) was used according to the manufacturer’s protocol. Specific probes for *C3* (Cat. Nr. 417848, ACDbio), *Gfap* (Cat. Nr. 313211-C2, ACDbio), and *Aldh1l1* (Cat. Nr. 405891-C2, ACDbio) were used. The astrocyte-specific probes *Gfap* and *Aldh1l1* were mixed 1:1 (astrocyte probe mix). Probes were detected using Opal dye 570 (FP1488001KT, Akoya Biosciences) for *C3* and Opal dye 520 (FP1487001KT) for the astrocyte probe mix. In addition, after RNAscope detection of C3 some slides were co-labelled with an anti-GFAP antibody (1:400, mab360, Millipore) according to the immunohistochemistry methods.

### Image Analysis

All image analysis was performed blinded. Images were either scanned on a Panoramic 250 Flash II slide scanner (3DHISTECH) or acquired with a DM6000 B microscope (Leica Microsystems) or a laser scanning confocal microscope (Carl Zeiss LSM 710). The medial corpus callosum was used for all quantifications. MBP immunohistochemistry was quantified using ImageJ Software v1.47 (Rasband, W.S., National Institutes of Health, Bethesda, MD, USA, http://imagej.nih.gov/ij). A user-defined macro was used to quantify the MBP positive signal area within the corpus callosum that was above a defined signal intensity threshold as a percentage of the defined area. For the quantification of RNAscope in situ hybridization signals, only cells with more than three fluorescence probe spots were counted as positive for a given probe.

### Rotarod

Motor coordination and balance were tested in injured and healthy WT and TKO mice at 28 days post injury (28dpi) using the rotarod. The rotarod test was performed as in Renz et al. 2022 [15]. Briefly, on the first day, the mice were trained to remain on the wheel at 5rpm for five minutes. The next day, the rotational speed was continuously increased from 15 to 33 rotations per minute (rpm), alternating between forward and backward rotation over 5 minutes. The mean latency to fall across three trials was recorded.

### Novel object recognition

Recognition memory was assessed at 28 dpi using the novel object recognition task [29]. Briefly, the animals were habituated to the open field one day before the task. On the following day, mice were returned to the same open field containing two identical objects. Four hours later, the mice explored the open field again in the presence of one familiar object and one new object (novel object). The order of object presentation and the relative object positions were randomized between mice. Total object exploration time was limited to 20 seconds with a maximum of 10 minutes in the open field. The discrimination ratio was calculated as novel object interaction time/total object interaction time with both objects.

### Preparation of live brain cell suspensions and FACS isolation of astrocytes

Live single-cell suspensions from a single healthy and injured mouse brain 1 day after injury (1 dpi) were prepared as follows: Cortices and underlying white matter were enzymatically digested using papain and then mechanically dissociated to yield a single cell suspension according to the immunopanning protocol of [30]. The cells were resuspended in Dulbecco’s PBS (DPBS) containing 0.02% BSA and 125 U/ml DNase, filtered through a nitex mesh filter (Tetko HC3-20) and supplemented with 1 μg/ml propidium iodide (PI) for fluorescence-activated cell sorting (FACS) [23]. Subsequently, astrocytes were sorted at 4°C using BD FACS ARIA III for GFP fluorescence in the absence of PI to select live astrocytes at the Flow Cytometry and Cell Sorting Facility (FCCS) of the Department for BioMedical Research, University of Bern, Switzerland.

### mRNA isolation and microfluidic qRT-PCR

Total RNA was isolated from FACS-purified astrocytes from a single mouse brain using the QIAshredder and the All-prep DNA/RNA/protein Mini Kit according to the manufacturer’s protocol (Qiagen). Total RNA concentration was measured using a NanoVue PlusTM spectrophotometer (Biochrom). Microfluidic qRT-PCR was performed using the Biomark HD System from Fluidigm according to the manufacturer’s protocol using the Delta Gene™ Assay and a 96.96 Dynamic Array™ IFC for Gene Expression. Briefly, cDNA was prepared using the reverse transcriptome master mix (PN 100-6297) followed by preamplification with the Preamp Master Mix (PN 100-5580) using the GE Fast 96×96 PCR + Melt v2 thermal cycling protocol according to the manufacturer’s instructions (Standard Biotools). The 96.96 Dynamic Array™ IFC was primed (including diluter cDNA and primer mastermix using the Biomark HX system IFC Controller. The loaded chip was processed in the Biomark HD Real-Time PCR System (Standard Biotools) and cycled as follows: 10min at 95°C followed by 40 cycles of 95°C for 15s, 60°C for 30s and 72°C for 30s. Data were collected and quantified using BioMark Data Collection Software 2.1.1 build 20090519.0926 from Standard Biotools. Expression data for each sample were normalized to the sample mean of *Rplp0* and *Aldh1l1* expression (Supplementary Table 4). Fold change for each gene was calculated relative to the mean expression of the corresponding gene in uninjured samples.

### Human post-mortem neonatal brain specimens

Human postmortem fixed-frozen brain tissue samples obtained from University of California, San Francisco’s Pediatric Neuropathology Research Laboratory were used for this study (Supplementary Table 1). All human tissue was collected in accordance with guidelines established by the University of California, San Francisco Committee on Human Research (H11170-19113-07) following the provision of informed consent. The procurement of de-identified postmortem human brain tissues was approved by the Institutional Review Board at the University of California San Francisco (#12-08643).Human postmortem tissue was fixed with 4% PFA and subsequently cryopreserved sequentially in 10%, 20%, and 30% sucrose. Frozen tissue was then cut into 14 μm sections for further examination. White matter injury was diagnosed by an experienced neuropathologist by the presence of astrogliosis and macrophage infiltration. Tissue from infants that died as a consequence of VATER malformations was used as a control. WMI and control cases were matched as closely as possible based on gestational age at the time of birth, postnatal age at the time of death, and brain region.

### Staining of human post-mortem tissue

In situ hybridization combined with immunohistochemistry was performed on fixed-frozen post-mortem brain sections using the the RNAscope^®^ Multiplex Fluorescent Reagent Kit v2 (Cat. Nr. 323135, ACDbio). Briefly, sections were prepared as follows: washed in PBS for 5 min, baked at 60°C for 30 min, fixed in 4% PFA on ice, dehydrated in 50%, 70% and 100% ethanol for 5 min at RT and then air dried. After hydrogen peroxide treatment for 10 min at RT, the target retrieval was performed for 5 min at 95°C. Subsequently, sections were treated with Protease III for 10 min at 40°C. The hybridization using probe *C3* (Cat. Nr. 430701-C3, ACDbio) and all further amplification steps were performed according to the manufacturer’s protocol. After the detection step using Opal dye 570 (SKU FP1488001KT, Akoya Biosciences), sections were washed in RNAscope^®^ wash buffer and blocked in 5% horse serum, 0.2% Triton X-100 (Sigma Aldrich) in 1x PBS for 1h at RT. Subsequently, sections were incubated overnight at 4°C with the primary anti-GFAP antibody (1:400, G3893, Sigma Aldrich). Next, sections were incubated with the secondary antibody (Alexa fluor^®^ 488-conjugated, Thermo Fisher Scientific) for 1h at RT and counterstained with DAPI for 30 seconds. All tissue sections were scanned using the Panoramic 250 Flash II slide scanner for quantification. All data analysis was performed blinded. The percentage of GFAP positive astrocytes labeled with *C3* was counted in 3 - 6 evenly spaced circular regions (0.5 mm^2^) placed across the white matter region of interest. GFAP expressing cells showing more than five *C3* RNAScope^®^ fluorescence probe spots were counted as positive.

### Statistical Analysis

The sample size for all our analyses was N < 30 (Supplementary Table 2). Consequently, we were unable to reliably test for a normal distribution of the data. Therefore, throughout the manuscript, we used the Mann-Whitney test for pairwise comparisons of continuous variables. Significance was determined at *p* < 0.05.

## Author Contributions

PR, DS, AS, and ABL conceptualized the project. PR and ABL designed experiments and analyzed the data with support of AS and DS. PR and VH performed the experiments. VT provided technical assistance. PR and ABL wrote the paper. All authors reviewed, corrected, and edited the manuscript. DR and MC provided human tissue and guided design of human tissue experiments. EH performed histopathological analysis of human tissue. SAL provided the TKO mice. DS and ABL obtained project funding.

## Institutional Review Board Statement

The animal study protocol was approved by the Ethics Committee of Canton Bern (reference number: BE19/85). The procurement of de-identified postmortem human brain tissues was approved by the Institutional Review Board at the University of California San Francisco (#12-08643).

## Data Availability Statement

The data presented in this study are available on request from the corresponding author.

## Acknowledgments

We express our appreciation to Marialuigia Giovannini-Spinelli, Marianne Jörger-Messerli, and Sophie Cottagnoud for helpful scientific input and experimental support. In addition, we are grateful to Smita Saxena, Federica Pilotto, and Mert Duman, Department of Neurology, Inselspital, University Hospital, and Department for BioMedical Research, University of Bern, Bern, Switzerland, for guidance with the rotarod test. D.H.R acknowledges the Dr Miriam and Sheldon G Adelson Medical Research Foundation, European Research Council Advanced Grant (no 789054) and NIH (P01NS08351) for support. M.C acknowledges funding from NIH (R00NS117804). SAL acknowledges the generous support of Paul Slavik and other anonymous donors.

