## Supplementary Tables for "Silencing neuroinflammatory reactive astrocyte activating factors ameliorates disease outcomes in perinatal white matter injury"

Supplementary Table 1. White matter injury and control cases of human postmortem brain tissue

| Case | Case number | Gestational age at birth | Postnatal age at time of death | Cause of death |
|---|---|---|---|---|
| WMI Case 1 | UCSF 2013-006 | 32 week 4 days | 12 days | Complications of prematurity |
| Control Case 1 | UCSF 2012-003 | 36 week 6 days | 10 days | VATER malformation |
| WMI Case 2 | UCSF2016-003 | 25 week | 8 months | Pre-term birth complications |
| Control Case 2 | UCSF2011-010 | 31 week | 7 months | Pnemonia, VATER malformation |

Supplementary Table 2. Summary of experimental cohorts

| Mouse Nr | | Strain | | Treatment | | IHC | | ISH + IHC |
|---|---|---|---|---|---|---|---|---|
| | | | | | | MBP | C3 + Aldh1l1 + Gfap | C3 + GFAP |
| **M83** | 16 | WT | P11 | 2 mg/kg NaCl | Normoxia 25 MIN | x | x | |
| **M84** | 16 | WT | P11 | 2 mg/kg NaCl | Normoxia 25 MIN | x | x | |
| **M85** | 16 | WT | P11 | 2 mg/kg NaCl | Normoxia 25 MIN | x | x | |
| **M86** | 16 | WT | P11 | 2 mg/kg LPS | 25 MIN @ 8% O2 | x | x | |
| **M87** | 16 | WT | P11 | 2 mg/kg LPS | 25 MIN @ 8% O2 | x | x | |
| **M88** | 16 | WT | P11 | 2 mg/kg LPS | 25 MIN @ 8% O2 | x | x | |
| **M89** | 16 | TKO | P11 | 2 mg/kg NaCl | Normoxia 25 MIN | x | | |
| **M90** | 16 | TKO | P11 | 2 mg/kg NaCl | Normoxia 25 MIN | x | | |
| **M91** | 16 | TKO | P11 | 2 mg/kg NaCl | Normoxia 25 MIN | x | | |
| **M92** | 16 | TKO | P11 | 2 mg/kg LPS | 25 MIN @ 8% O2 | x | | |
| **M93** | 16 | TKO | P11 | 2 mg/kg LPS | 25 MIN @ 8% O2 | x | | |
| **M94** | 16 | TKO | P11 | 2 mg/kg LPS | 25 MIN @ 8% O2 | x | | |
| **M95** | 16 | TKO | P11 | 2 mg/kg LPS | 25 MIN @ 8% O2 | x | | |
| **M96** | 16 | TKO | P11 | 2 mg/kg LPS | 25 MIN @ 8% O2 | x | | |
| **M99** | 16 | WT | P11 | 2 mg/kg NaCl | Normoxia 25 MIN | x | x | |
| **M100** | 16 | WT | P11 | 2 mg/kg NaCl | Normoxia 25 MIN | x | x | |
| **M101** | 16 | WT | P11 | 2 mg/kg NaCl | Normoxia 25 MIN | x | x | |
| **M102** | 17 | TKO | P11 | 2 mg/kg NaCl | Normoxia 25 MIN | x | | |
| **M103** | 17 | TKO | P11 | 2 mg/kg NaCl | Normoxia 25 MIN | x | | |
| **M104** | 17 | TKO | P11 | 2 mg/kg LPS | 25 MIN @ 8% O2 | x | | |
| **M105** | 17 | TKO | P11 | 2 mg/kg LPS | 25 MIN @ 8% O2 | x | | |
| **M110** | 21 | TKO | P11 | 2 mg/kg NaCl | Normoxia 25 MIN | x | | |
| **M111** | 21 | TKO | P11 | 2 mg/kg NaCl | Normoxia 25 MIN | x | | |
| **M112** | 21 | TKO | P11 | 2 mg/kg LPS | 25 MIN @ 8% O2 | x | | |
| **M113** | 21 | TKO | P11 | 2 mg/kg LPS | 25 MIN @ 8% O2 | x | | |
| **M114** | 21 | TKO | P11 | 2 mg/kg LPS | 25 MIN @ 8% O2 | x | | |
| **M119** | 21 | WT | P11 | 2 mg/kg NaCl | Normoxia 25 MIN | x | x | |
| **M121** | 21 | WT | P11 | 2 mg/kg LPS | 25 MIN @ 8% O2 | x | x | |
| **M122** | 21 | WT | P11 | 2 mg/kg LPS | 25 MIN @ 8% O2 | x | x | |
| **M123** | 21 | WT | P11 | 2 mg/kg LPS | 25 MIN @ 8% O2 | x | x | |
| **M135** | 23 | WT | P3 | 2 mg/kg NaCl | Normoxia 25 MIN | | x | x |
| **M136** | 23 | WT | P3 | 2 mg/kg NaCl | Normoxia 25 MIN | | x | x |
| **M137** | 23 | WT | P3 | 2 mg/kg LPS | 25 MIN @ 8% O2 | | x | x |
| **M138** | 23 | WT | P3 | 2 mg/kg LPS | 25 MIN @ 8% O2 | | x | x |
| **M139** | 23 | WT | P3 | 2 mg/kg LPS | 25 MIN @ 8% O2 | | x | x |
| **M140** | 23 | WT | P4 | 2 mg/kg NaCl | Normoxia 25 MIN | | x | |
| **M141** | 23 | WT | P4 | 2 mg/kg NaCl | Normoxia 25 MIN | | x | |
| **M142** | 23 | WT | P4 | 2 mg/kg LPS | 25 MIN @ 8% O2 | | x | |
| **M143** | 23 | WT | P4 | 2 mg/kg LPS | 25 MIN @ 8% O2 | | x | |
| **M144** | 23 | WT | P11 | 2 mg/kg NaCl | Normoxia 25 MIN | x | x | |
| **M145** | 23 | WT | P11 | 2 mg/kg LPS | 25 MIN @ 8% O2 | x | x | |
| **M146** | 24 | WT | P3 | 2 mg/kg NaCl | Normoxia 25 MIN | | x | |
| **M147** | 24 | TKO | P3 | 2 mg/kg NaCl | Normoxia 25 MIN | | x | |
| **M148** | 24 | TKO | P3 | 2 mg/kg NaCl | Normoxia 25 MIN | | x | |
| **M149** | 24 | TKO | P3 | 2 mg/kg NaCl | Normoxia 25 MIN | | x | |
| **M150** | 24 | TKO | P3 | 2 mg/kg LPS | 25 MIN @ 8% O2 | | x | |
| **M151** | 24 | TKO | P3 | 2 mg/kg LPS | 25 MIN @ 8% O2 | | x | |
| **M152** | 24 | TKO | P3 | 2 mg/kg LPS | 25 MIN @ 8% O2 | | x | |
| **M153** | 24 | TKO | P3 | 2 mg/kg LPS | 25 MIN @ 8% O2 | | x | |
| **M154** | 24 | WT | P4 | 2 mg/kg NaCl | Normoxia 25 MIN | | x | |
| **M155** | 24 | WT | P4 | 2 mg/kg NaCl | Normoxia 25 MIN | | x | |
| **M156** | 24 | WT | P4 | 2 mg/kg NaCl | Normoxia 25 MIN | | x | |
| **M157** | 24 | WT | P4 | 2 mg/kg LPS | 25 MIN @ 8% O2 | | x | |
| **M158** | 24 | WT | P11 | 2 mg/kg NaCl | Normoxia 25 MIN | x | x | |
| **M159** | 24 | WT | P11 | 2 mg/kg NaCl | Normoxia 25 MIN | x | x | |
| **M160** | 25 | WT | P3 | 2 mg/kg NaCl | Normoxia 25 MIN | | x | x |
| **M161** | 25 | WT | P3 | 2 mg/kg LPS | 25 MIN @ 8% O2 | | x | x |
| **M162** | 25 | WT | P3 | 2 mg/kg NaCl | Normoxia 25 MIN | | x | x |
| **M163** | 25 | WT | P3 | 2 mg/kg LPS | 25 MIN @ 8% O2 | | x | |
| **M165** | 25 | WT | P3 | 2 mg/kg LPS | 25 MIN @ 8% O2 | | x | x |
| **M166** | 25 | WT | P3 | 2 mg/kg LPS | 25 MIN @ 8% O2 | | x | |
| **M167** | 25 | WT | P4 | 2 mg/kg NaCl | Normoxia 25 MIN | | x | |
| **M168** | 25 | WT | P4 | 2 mg/kg LPS | 25 MIN @ 8% O2 | | x | |
| **M169** | 25 | WT | P4 | 2 mg/kg LPS | 25 MIN @ 8% O2 | | x | |
| **M170** | 25 | WT | P4 | 2 mg/kg LPS | 25 MIN @ 8% O2 | | x | |
| **M171** | 25 | WT | P4 | 2 mg/kg LPS | 25 MIN @ 8% O2 | | x | |

Supplementary Table 3. P-values from microfluidic qRT-PCR

| Gene | P value | |
|---|---|---|
| | WT | TKO |
| Aspg | 0.003126 | 0.304204 |
| CD44 | 0.014746 | 0.077522 |
| CP | 0.006454 | 0.766649 |
| Cxcl10 | 0.029031 | 0.223496 |
| Hsbp1 | 0.808929 | 0.842864 |
| Lcn2 | 0.045947 | 0.070955 |
| Osmr | 0.059169 | 0.742079 |
| S1pr3 | 0.136673 | 0.650482 |
| Serpina3n | 0.017779 | 0.268463 |
| Steap4 | 0.690531 | 0.166046 |
| Gfap | 0.005976 | 0.210239 |
| Vim | 0.50724 | 0.78866 |
| Amigo2 | 0.766972 | 0.249717 |
| Fbln5 | 0.030115 | 0.867009 |
| Fkbp5 | 0.061412 | 0.356578 |
| Gbp2 | 0.020823 | 0.255845 |
| H2-D1 | 0.001747 | 0.399951 |
| H2-T23 | 0.003889 | 0.017392 |
| Ligp1 | 0.004609 | 0.056511 |
| Psmb8 | 0.000024 | 0.107755 |
| Serping1 | 0.060149 | 0.113004 |
| Srgn | 0.322486 | 0.342287 |
| Ugt1a | 0.421246 | 0.029549 |
| Ggta1 | 0.08476 | 0.080928 |
| C3 | 0.044035 | 0.297252 |
| B3gnt5 | 0.024793 | 0.876718 |
| CD109 | 0.359796 | 0.776009 |
| CD14 | 0.002171 | 0.781241 |
| Clcf1 | 0.173174 | 0.000239 |
| Emp1 | 0.028701 | 0.537093 |
| Ptgs2 | 0.02151 | 0.950525 |
| Ptx3 | 0.997839 | 0.835317 |
| S100a10 | 0.21683 | 0.432691 |
| Slc10a6 | 0.764436 | 0.963682 |
| Sphk1 | 0.621573 | 0.445153 |
| Tgm1 | 0.076939 | 0.154919 |
| Tm4sf1 | 0.429855 | 0.989504 |

Supplementary Table 4. Summary of primers used for microfluidic qRT-PCR

| Fluidigm Delta Gene Assays Target | | |
|---|---|---|
| **Name** | **Forward Primer Sequence** | **Reverse Primer Sequence** |
| Rplp0 | AGATTCGGGATATGCTGTTGGC | TCGGGTCCTAGACCAGTGTTC |
| Aldh1l1 | GCAGGTACTTCTGGGTTGCT | GGAAGGCACCCAAGGTCAAA |
| Gapdh | AAGAGGGATGCTGCCCTTAC | TACGGCCAAATCCGTTCACA |
| Aspg | GCTGCTGGCCATTTACACTG | GTGGGCCTGTGCATACTCTT |
| Cd44 | ACCTTGGCCACCACTCCTAA | GCAGTAGGCTGAAGGGTTGT |
| Cp | TGTGATGGGAATGGGCAATGA | AGTGTATAGAGGATGTTCCAGGTCA |
| Cxcl10 | CCCACGTGTTGAGATCATTG | CACTGGGTAAAGGGGAGTGA |
| Gfap | AGAAAGGTTGAATCGCTGGA | CGGCGATAGTCGTTAGCTTC |
| Hsbp1 | GACATGAGCAGTCGGATTGA | GGATGGGGTGTAGGGGTACT |
| Lcn2 | CCAGTTCGCCATGGTATTTT | CACACTCACCACCCATTCAG |
| Osmr | GTGAAGGACCCAAAGCATGT | GCCTAATACCTGGTGCGTGT |
| S1pr3 | AAGCCTAGCGGGAGAGAAAC | TCAGGGAACAATTGGGAGAG |
| Serpina3n | CCTGGAGGATGTCCTTTCAA | TTATCAGGAAAGGCCGATTG |
| Steap4 | CCCGAATCGTGTCTTTCCTA | GGCCTGAGTAATGGTTGCAT |
| Vim | AGACCAGAGATGGACAGGTGA | TTGCGCTCCTGAAAAACTGC |
| Amigo2 | GAGGCGACCATAATGTCGTT | GCATCCAACAGTCCGATTCT |
| Fbln5 | CTTCAGATGCAAGCAACAA | AGGCAGTGTCAGAGGCCTTA |
| Fkbp5 | TATGCTTATGGCTCGGCTGG | CAGCCTTCCAGGTGGACTTT |
| Gbp2 | GGGGTCACTGTCTGACCACT | GGGAAACCTGGGATGAGATT |
| Ggta1 | GTGAACAGCATGAGGGGTTT | GTTTTGTTGCCTCTGGGTGT |
| H2-D1 | TCCGAGATTGTAAAGCGTGAAGA | ACAGGGCAGTGCAGGGATAG |
| H2-T23 | GGACCGCGAATGACATAGC | GCACCTCAGGGTGACTTCAT |
| Iigp1 | GGGGCAATAGCTCATTGGTA | ACCTCGAAGACATCCCCTTT |
| Psmb8 | CAGTCCTGAAGAGGCCTACG | CACTTTCACCCAACCGTCTT |
| Serping1 | ACAGCCCCCTCTGAATTCTT | GGATGCTCTCCAAGTTGCTC |
| Srgn | GCAAGGTTATCCTGCTCGGA | TGGGAGGGCCGATGTTATTG |
| Ugt1a | CCTATGGGTCACTTGCCACT | AAAACCATGTTGGGCATGAT |
| C3 | CGCAACGAACAGGTGGAGATCA | CTGGAAGTAGCGATTCTTGGCG |
| B3gnt5 | CGTGGGGCAATGAGAACTAT | CCCAGCTGAACTGAAGAAGG |
| Cd109 | CACAGTCGGGAGCCCTAAAG | GCAGCGATTTCGATGTCCAC |
| Cd14 | GGACTGATCTCAGCCCTCTG | GCTTCAGCCCAGTGAAAGAC |
| Clcf1 | CTTCAATCCTCCTCGACTGG | TACGTCGGAGTTCAGCTGTG |
| Emp1 | GAGACACTGGCCAGAAAAGC | TAAAAGGCAAGGGAATGCAC |
| Ptgs2 | GCTGTACAAGCAGTGGCAAA | CCCCAAAGATAGCATCTGGA |
| Ptx3 | AACAAGCTCTGTTGCCCATT | TCCCAAATGGAACATTGGAT |
| S100a10 | CCTCTGGCTGTGGACAAAAT | CTGCTCACAAGAAGCAGTGG |
| Slc10a6 | GCTTCGGTGGTATGATGCTT | CCACAGGCTTTTCTGGTGAT |
| Sphk1 | GATGCATGAGGTGGTGAATG | TGCTCGTACCCAGCATAGTG |
| Tgm1 | CTGTTGGTCCCGTCCCAAA | GGACCTTCCATTGTGCCTGG |
| Tm4sf1 | GCCCAAGCATATTGTGGAGT | AGGGTAGGATGTGGCACAAG |
