## Supplementary Figures for "Silencing neuroinflammatory reactive astrocyte activating factors ameliorates disease outcomes in perinatal white matter injury"

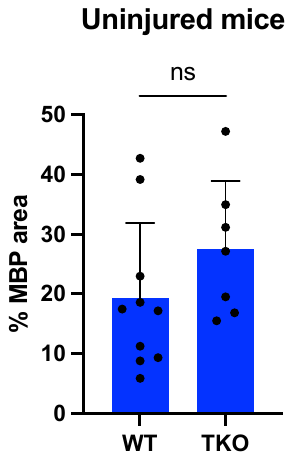

**Supplementary Figure 1.** Quantification of myelination defects. Quantification of MBP IHC signal in the medial corpus callosum at 9 dpi in uninjured brains of wild type (WT) C57Bl/6 (n = 10) and *Tnf, Il1a, C1qa* triple knockout (TKO, n = 7) mice. Data are presented as mean ± SD. ns not significant.

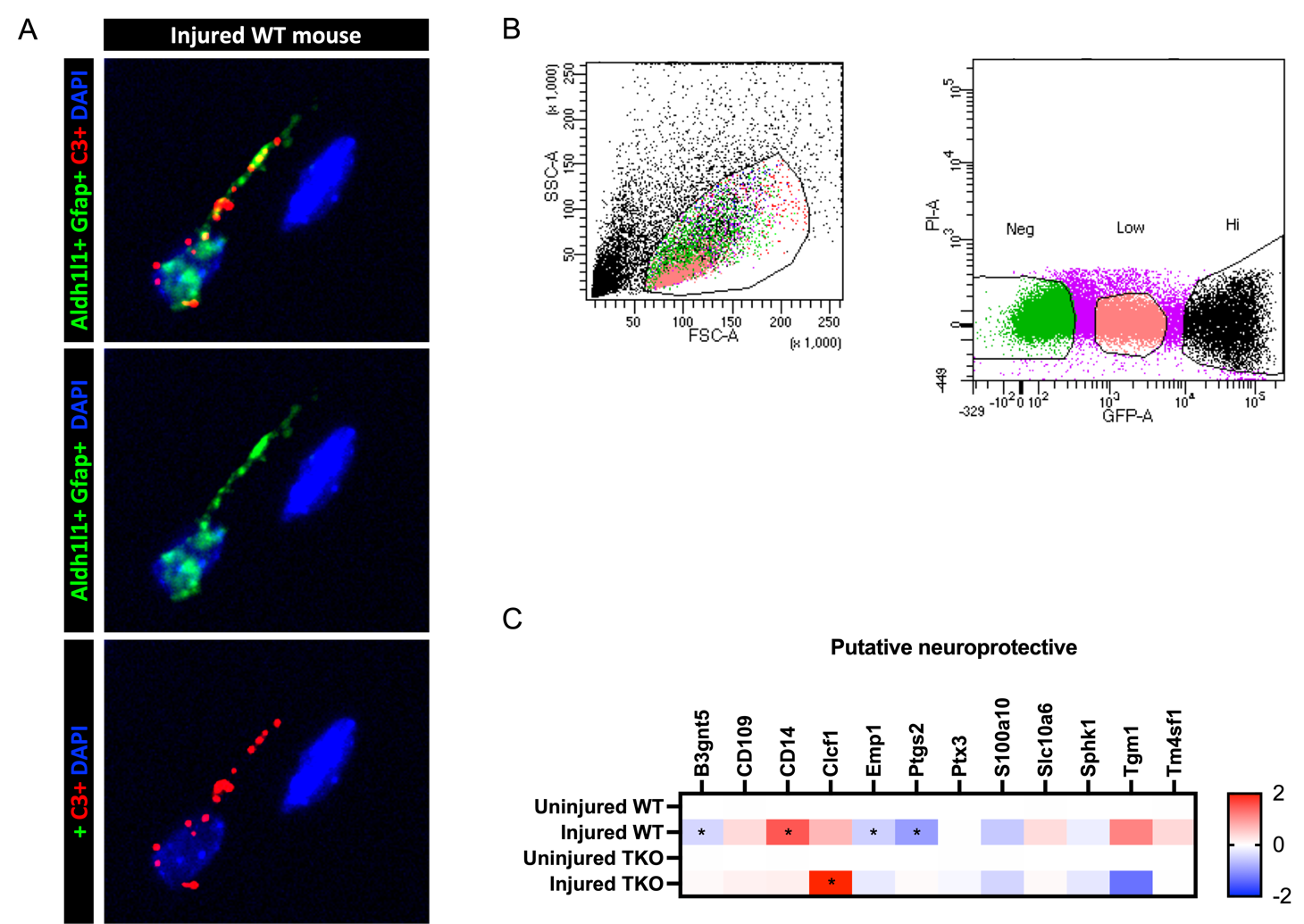

**Supplementary Figure 2.** (A) In situ hybridization for inflammatory reactive astrocyte marker *C3* (red) and astrocyte markers *Aldh1l1* and *Gfap* (probe mix, green) in the mouse corpus callosum of an injured WT brain. Scale bar: 1000 μm. (B) FACS plots indicating the side scatter (SSC), forward scatter (FSC), and subsequent green fluorescent protein (GFP) cutoffs used to select a cell population (GFP Hi) for qRT-PCR analysis. (C) Microfluidic qRT-PCR for a panel of genes reported to be representative of a neuroprotective reactive astrocyte state. Heatmap compares the mean fold change (FC) of neuroprotective gene transcripts from astrocytes isolated from the cortex and white matter of uninjured and injured wild type (WT) C57Bl/6 (n = 6) or *Tnf, Il1a, C1qa* triple knockout (TKO, n = 4) mice.
